# Monkey Features Location Identification Using Convolutional Neural Networks

**DOI:** 10.1101/377895

**Authors:** Rollyn Labuguen (P), Vishal Gaurav, Salvador Negrete Blanco, Jumpei Matsumoto, Kenichi Inoue, Tomohiro Shibata

## Abstract

Understanding animal behavior in its natural habitat is a challenging task. One of the primary step for analyzing animal behavior is feature detection. In this study, we propose the use of deep convolutional neural network (CNN) to locate monkey features from raw RGB images of monkey in its natural environment. We train the model to identify features such as the nose and shoulders of the monkey at about 0.01 model loss.

## 1 Introduction

Attempts on automated body-parts tracking on insects and mammals have often relied in attaching markers in the animal, and then a software keeps track of these markers [4]. However, the markers can affect the animals natural behavior. For this reason, low-cost marker-less motion capture systems for animal tracking are developed and introduced in [2] and [3] using depth sensors. Recent studies also utilized convolutional neural networks to do marker-less 2D animal pose estimation such as [5]; however, they have focusd more in laboratory-controlled setups and their results are limited to the fruit fly and mouse. The specific targets of the previous works are typically insects and rodents; few have been dedicated for primates and monkeys. On this note, MONET [6] has used multiview image streams for monkey pose detection with a semi-supervised learning framework, but it has not use any deep learning methods, and only relied upon time and space consistency across image streams for annotation augmentation.

In contrast to traditional image segmentation and feature engineering, learning interaction and representation from data is the core of deep learning [1] application to real-world scenarios. For this study, we apply deep CNN as means to locate key feature points of monkey towards understanding its behavior in its natural habitat. The use of CNN forgoes the use of hand engineered features and allows the model to learn monkey features from data as training proceeds.

## 2 Proposed Methodology

Our method for finding monkey features is composed of convolutional layers for feature learning and then max pooling as depicted in Figure 2. For any given input image with corresponding vector that contains the monkey features’ 2D coordinate locations, specifically *{*nose; left shoulder; right shoulder*}*. The latent representation of the extracted feature map from input image *I_i_* is given by Eq. (1):

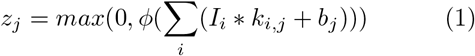

where *I_i_* and *z_j_* denotes *i^th^* input map and *j^th^* output map respectively and ∗ denotes the convolution operation. *k_i,j_ and b_j_* are kernel and bias. *ϕ* is Max Pooling operation to reduce the spatial resolution of feature maps. Rectified Linear Unit (ReLU) (*f*(*x*) = *max*(0*, x*) activation is used to induce non-linearity and faster convegence. We used Mean Squared Error (MSE) as loss function given by Eq. (2)

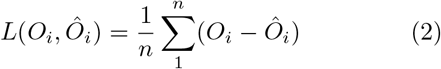

Adam optimizer is used to minimize the loss between predicted output *Ô_i_* and actual annotated *O_i_* feature location.

**Figure 1:**
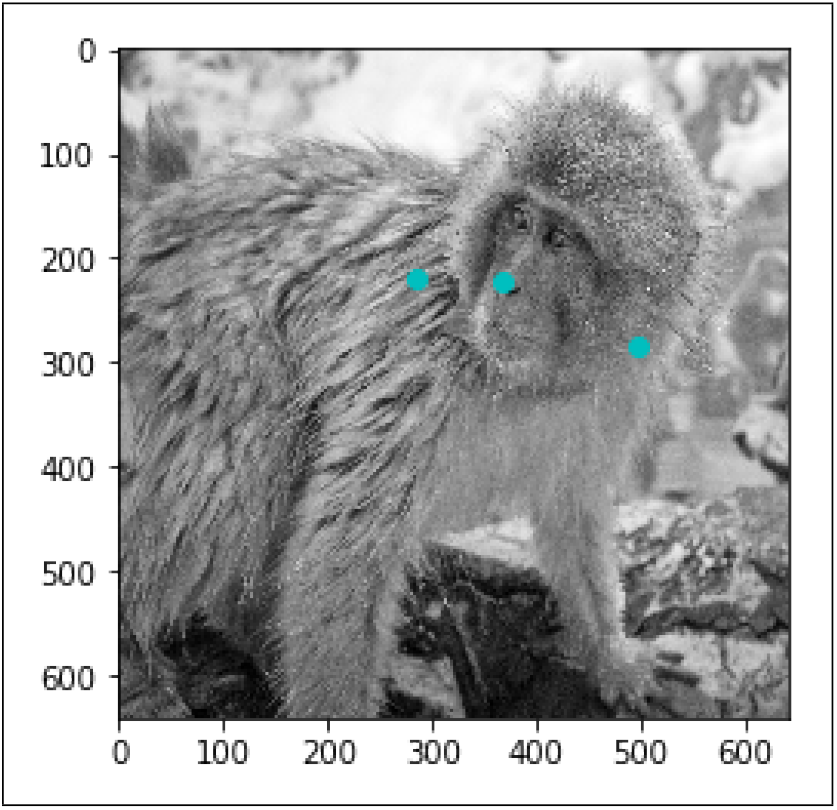
Example of monkey image used for training with keypoints annotation

**Figure 2:**
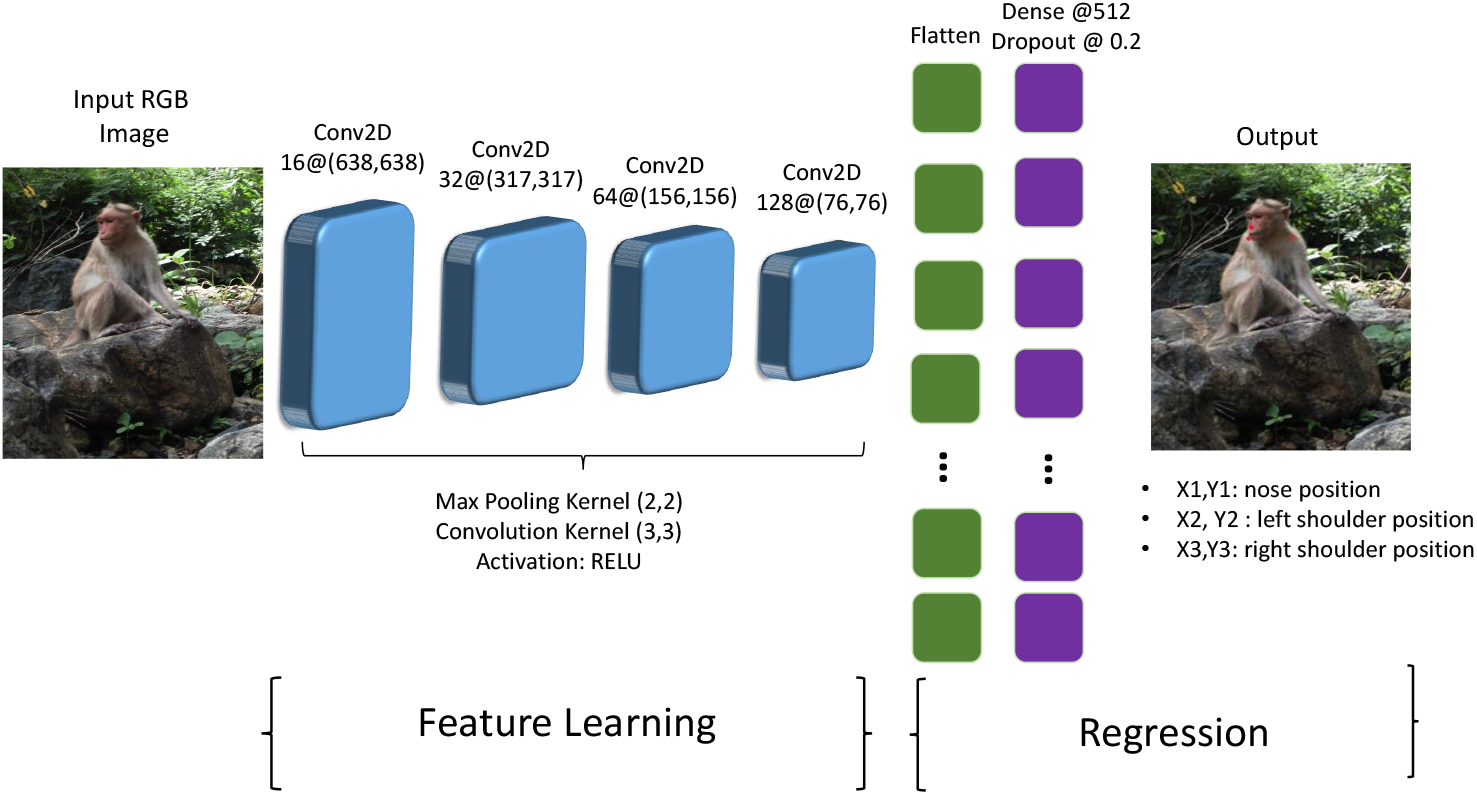
Pipeline for Monkey Feature Location Identification with Deep CNN of six hidden layers

### 2.1 Dataset Preparation

In collaboration with the Primate Research Institute, our dataset contains a total of 6000 high-resolution images of monkeys, which are manually annotated with the feature locations. We have divided the data set in three parts, 60% train, 20% validation and 20% test sets, respectively.

Each input data includes single monkey on the image and its pre-determined part annotations. To train and validate our CNN architecture for feature detection, we resized the images to (640 ∗ 640) pixels for faster computation and lesser memory allocation. To verify our proposed method, we only used the nose and shoulders of the monkey. Figure 1 shows example of monkey image in the dataset together with the annotations used.

### 2.2 Network Architecture

The CNN architecture shown in Figure 2 is used to locate and identify the essential monkey features. We designed a sequential CNN having four blocks of convolutional layers and max pooling, followed by a fully-connected layer with dropout of 20%. Filter sizes are indicated in each convolutional layer as follows: 16 for the first layer, 32 for the second layer, 64 for the third and 128 on the fourth. The expected output is normalized coordinates (*x_n_, y_n_*) of the *n* features in the monkey.

## 3 Results and Discussion

The deep learning CNN model is implemented in Python with Keras [7] libraries having Tensorflow [8] support and trained on GTX Titan X Graphics Processing Unit (GPU). The training time for our deep learning model was *≈* 30 minutes for 50 epochs. The validation and training loss of model is depicted in Figure 3. The training loss is significantly reduced after 30 epochs. Validation loss is consistent throughout the training, which implies that the network architecture must still be improved to predict the key feature points of the monkey.

**Figure 3:**
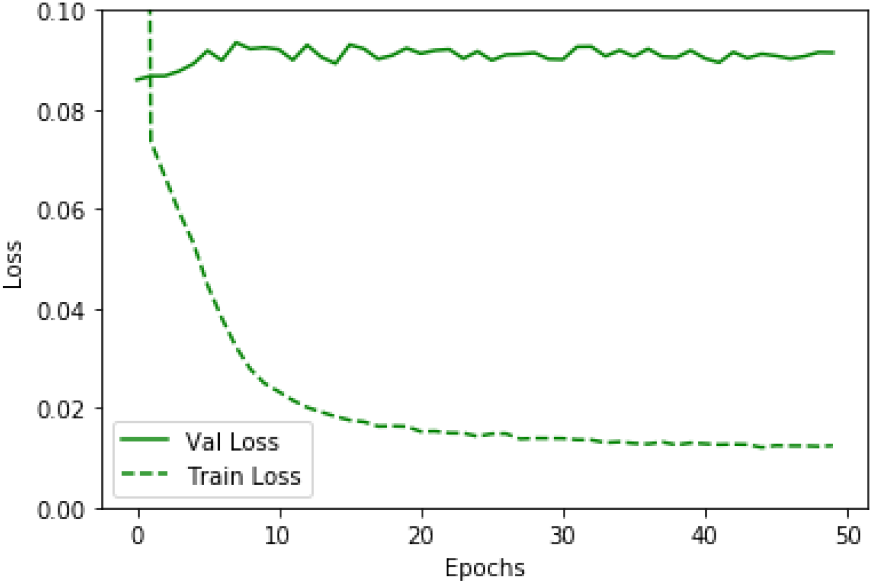
Train and Validation Loss at almost 0.01 and 0.09 respectively

Our CNN model is able to estimate the body part positions on the monkey regardless of the background change and its various poses. Figure 4 depicts the prediction of monkey features on unseen images. However, there are some errors in prediction of feature point for estimating monkey feature location. The predicted points are a bit shifted as depicted in the last row of Figure 4. While there are incorrect estimations, the predicted feature points are still located in the same area as the ground truth.

**Figure 4:**
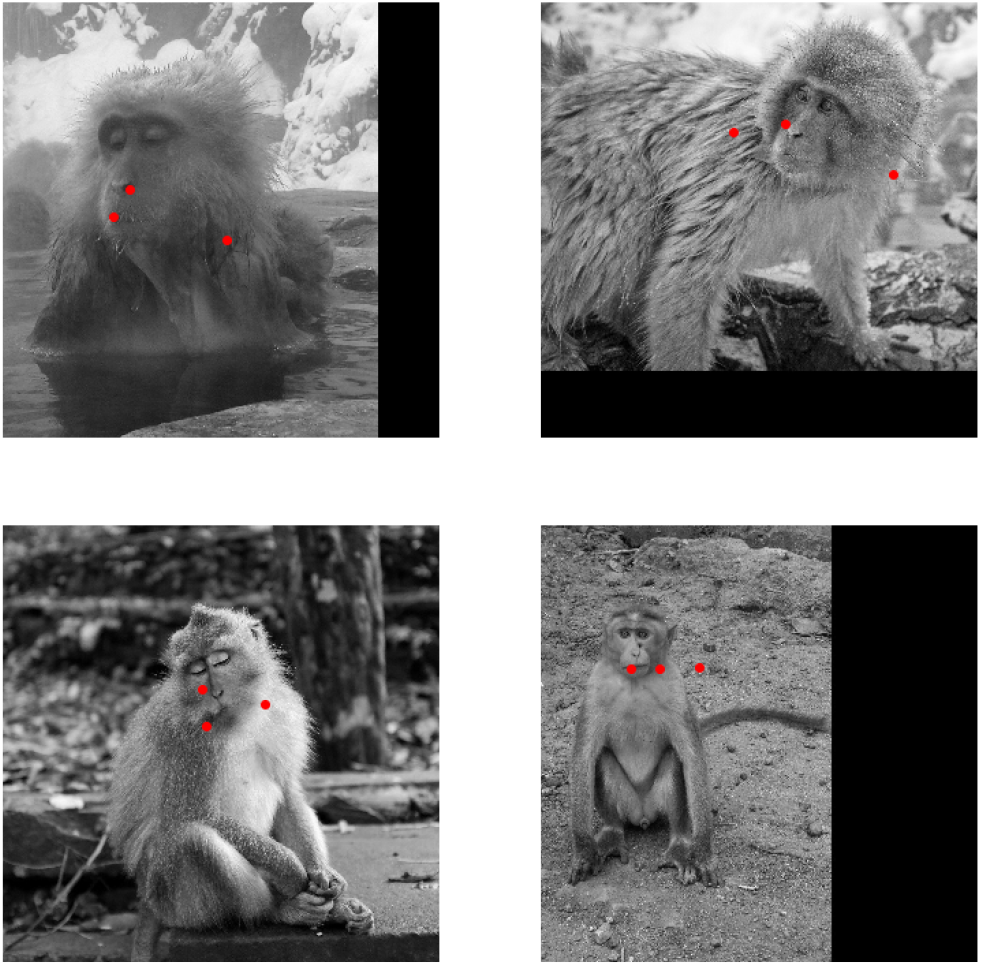
Feature extraction results (shown in red) of nose, left and right shoulders of the monkey

Figure 5 depicts the activation map on the final layer of the convolution process. As we go deeper, the activation becomes increasingly sparse and less visually interpretable. The network begin to encode the higher level features like “monkey face”, “monkey pose”, etc. In the first layer almost all filters are activated by the input image. But in subsequent layers, many filters are blank. This shows that the pattern encoded in those filters are not found in the input image. The activation maps on each convolutional layers demonstrate the inferred features of the network, such as the shape information of the monkey as well as the background. Most of the activations on the final convolutional layer “conv2d 4” have detected higher-level concepts and highlighted the pixel features on the face and body parts of the monkey as shown in Figure 5.

**Figure 5:**
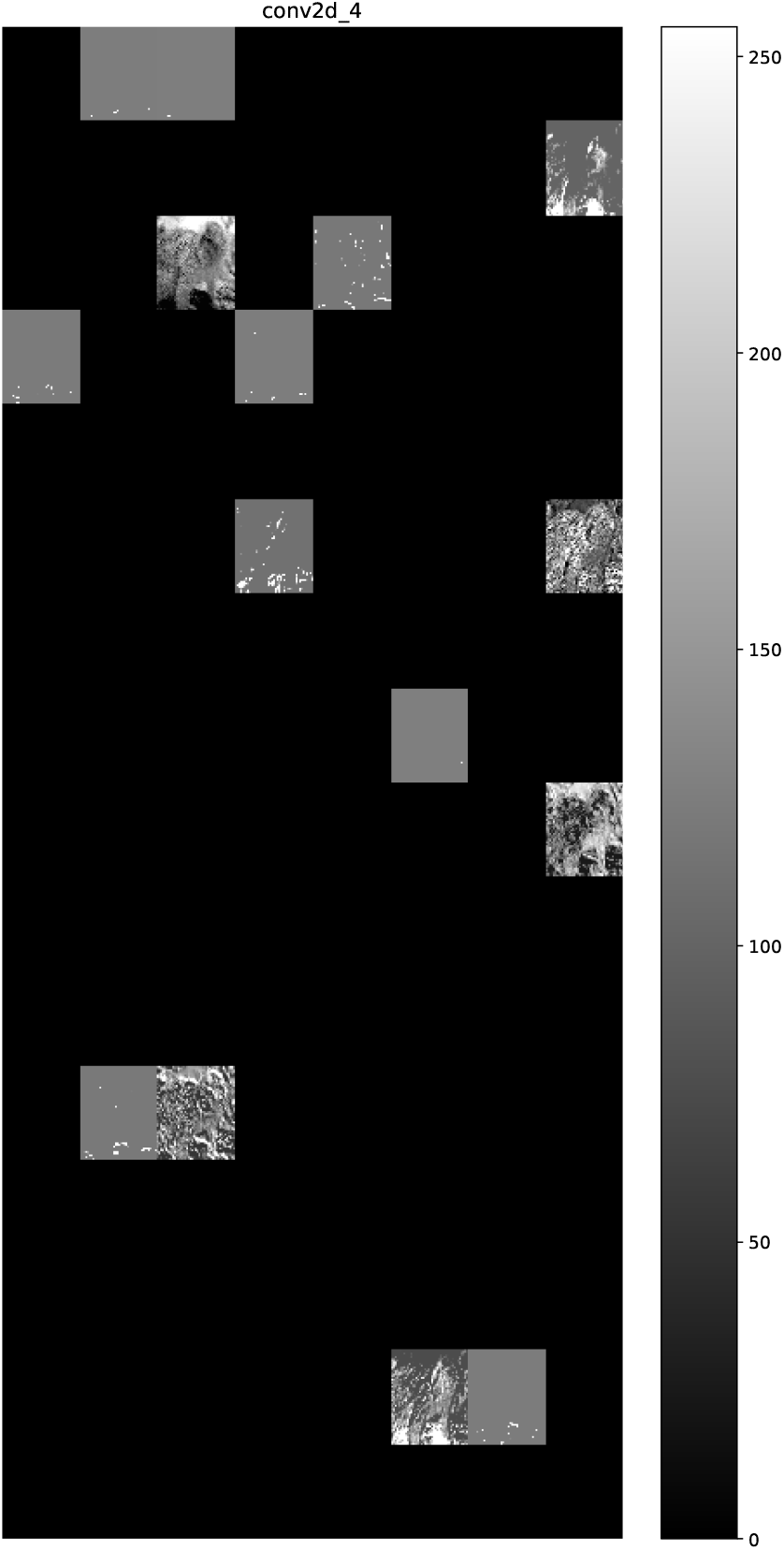
Final layer activation map showing the features encoded within the filters.

## 4 Conclusion and Future Work

In this study, we proposed a deep learning framework for predicting feature points in monkey images as an initial step towards behavior analysis of monkeys in its natural habitat. Specifically, we used a Convolutional Neural Network (CNN) regression model to predict the key features in monkey images. The trained model loss goes into plateau after several iterations at a very minimal value as depicted in Figure 3. From the activation maps of the CNN in Figure 5, we deduced that the model was able to discriminate between monkey features and the environment.

Furthermore, we want to incorporate more feature points for prediction and estimation of monkey pose. We plan to tune more on the network architecture varying the kernel size of the convolutional layers for prediction of monkey features. Nevertheless, this study has demonstrated the strength of using deep neural networks to identify and detect feature locations of essential body parts on primates.

## Acknowledgement

Authors would like to acknowledge the support given by Grant-in-Aid for Scientific Research from Japan Society for the Promotion of Science (No. 16H06534).

